# ELOVL2 is required for robust visual function in zebrafish

**DOI:** 10.1101/2020.10.13.336974

**Authors:** Manish Dasyani, Fangyuan Gao, Qianlan Xu, Donald Van Fossan, Ellis Zhang, Antonio F. M. Pinto, Alan Saghatelian, Dorota Skowronska-Krawczyk, Daniel L. Chao

**Affiliations:** Shiley Eye Institute, Viterbi Family Department of Ophthalmology, University of California San Diego; Department of Biophysics and Physiology, Center for Translational Vision Research, Gavin Hebert Eye Institute, University of California Irvine, Irvine, CA, USA; The Salk Institute for Biological Studies, Clayton Foundation Laboratories for Peptide Biology, 10010 N. Torrey Pines Rd, La Jolla, CA, 92037

## Abstract

Omega-3 and omega-6 polyunsaturated fatty acids (PUFAs) play critical roles in membrane stability and cell signaling within the retina. *Elovl2*, an elongase involved in synthesis of long chain polyunsaturated fatty acids (LC-PUFAs), has recently been implicated in regulating aging in the mammalian retina. In this work, we characterize the expression and function of *elovl2* in retina development in embryonic zebrafish. Whole mount *in situ* hybridization shows *elovl2* is expressed in the Müller glia in embryonic and adult zebrafish. Lipidomics analysis of *elovl2* crispants whole embryos at day 2 and eyes at day 7 demonstrated significant changes in lipids composition, especially on the level of lipids containing docosahexaenoic acid (DHA). Histological analysis of zebrafish lacking *elovl2* revealed increased retinal thickness compared to controls at day 7 without gross disruptions of retinal architecture. Finally, *elovl2* crispants showed differences in the visual motor reflex light off (VMR-OFF) at day 7 compared to controls. In sum, inactivation of *elovl2* in zebrafish embryos caused changes in lipid composition and in visual behavior further confirming the important role of LC-PUFAs in healthy vision.

## INTRODUCTION

Omega-3 and omega-6 polyunsaturated fatty acids (PUFAs), are critical for diverse biological functions including membrane stability, cell signaling and metabolism particularly in the brain and retina[1, 2]. Synthesis of PUFAs begins with dietary intake of essential amino acids, linoleic acid and alpha linoleic acid, which then goes through a series of elongation and desaturation reactions to form longer chain omega-3 and omega-6 fatty acids, including arachidonic acid, eicosapentaenoic acid (EPA), and docosahexaenoic acid (DHA). DHA, the major polyunsaturated fatty acid in the brain and retina, is a critical component of photoreceptor outer segments necessary for photoreceptor function[1, 3, 4] while very long chain PUFAs (VLC-PUFAs) are believed to be indispensable in maintaining the curvature of the photoreceptor disk membrane. PUFAs have been implicated in eye diseases, as decrease of dietary intake of food rich in omega-3 fatty acids, such as fish, have been linked to higher risk of age related eye diseases such as macular degeneration in multiple epidemiologic studies [4–6].

*ELOVL2* (Elongation Of Very Long Chain Fatty Acids-Like 2) encodes an enzyme involved in the elongation of long-chain (C20 and C22) omega-3 and omega-6 polyunsaturated fatty acids (LC-PUFAs) [7]. In particular, *ELOVL2* elongates docosapentaenoic acid (DPA − 22:5n-3) to 24:5n-3, which in turn is the substrate for the formation of VLC-PUFAs as well as 22:6n-3, i.e.. DHA [8]. Interestingly, DNA methylation of the *Elovl2* regulatory element has been well established as an epigenetic biomarker of aging, as multiple studies have shown that the DNA methylation of the *ELOVL2* promoter across multiple human tissues highly correlates with chronological age [9, 10].

We have previously studied the role of *Elovl2* in the mammalian retina. We have implicated *Elovl2* as a critical molecular regulator of aging, as loss of *Elovl2* activity in mice accelerates anatomical and functional surrogates of aging in the retina. Additionally, we observed subRPE deposits which contained multiple proteins found in human drusen, a pathologic hallmark of age related macular degeneration [11]. Pharmacological demethylation of *Elovl2* can increase *Elovl2* gene expression and prevent the progression of the age-related decline of the electroretinogram response, a functional surrogate of aging [11].

Zebrafish (*Danio rerio*) represents an excellent model system to study retinal development and biology, given its small size, transparent embryo, and the availability of robust visual behavior assays [12]. The role of *elovl2* in zebrafish, particularly within the eye, is still poorly understood. Previous studies have demonstrated that zebrafish *elovl2* can elongate C18-C22 PUFAs in a heterologous yeast system, in contrast to human ELOVL2 which has been shown to only elongate C20 and C22 PUFAs [7, 13, 14]. In addition, zebrafish *elovl5*, which in humans is specific to elongation of C18 PUFAs, can also elongate C20 and C22 PUFAs to a lesser extent. In a recent study of *elovl2* and *elovl5* knockout zebrafish, it was observed that *elovl2* but not *elovl5* is required for conversion of C20 EPA to DPA, and thus synthesis of DHA [15]. Finally, zebrafish *elovl4* is also able to elongate LC-PUFAs in striking contrast to the mammalian ortholog [16].

Here, we present the investigation of the function of *elovl2* in the zebrafish eye. We characterized the expression of *elovl2* in embryonic and adult wildtype zebrafish retina. We created zebrafish lacking *elovl2* function through introduction of biallelic mutations in *elovl2* using CRISPR-Cas9 technology, termed “crispants” [17]. Using lipidomics, retinal morphology, and visual behavior we showed the unexpected expression pattern and role of *elovl2* in zebrafish eye.

## MATERIALS AND METHODS

### Zebrafish husbandry

Fish were maintained in the fish facility of the University of California, San Diego (UCSD) under a controlled 14/10 hour light cycle with recirculating aquarium system. Breeding and experimental procedures were approved by the Institute of Animal Care and Use Committee of the University of California, San Diego (S18067).

### Zebrafish crispant creation

Potential gRNA sequences for zebrafish *elovl2* were searched for using the Chop-chop algorithm (https://chopchop.cbu.uib.no). Two *elovl2* gRNA sequences were selected, GACAGCCTATTTGGAGAAAG in exon 2 and TTCCCAGGTAGATTGTTAGG in exon 3. The control gRNA used (TGAGTATTCGCATGCAACTA) does not target any known zebrafish nucleotide sequence.

GRNA oligonucleotides (Integrated DNA Technologies (IDT), Coraville Iowa) were synthesized by and were duplexed individually with tracrRNA (Integrated DNA Technologies (IDT) Coraville, Iowa). The injection mix containing 250 ng/μl of gRNA duplex complex (both gRNAs), 500 ng/μl rCas9 protein (PNA Bio CP01-20 Thousand Oaks, California) and duplex buffer (IDT) was injected into one cell stage zebrafish embryos. The embryos were maintained in a 28°C incubator. The level of DNA editing was determined through DNA Sanger sequencing (Genewiz La Jolla CA) and analysis using the ICE v2CRISPR Analysis Tool (Synthego (Redwood City, CA), found at https://www.synthego.com/products/bioinformatics/crispr-analysis)

### Zebrafish *in-situ* hybridization

Whole mount RNA *in situ* hybridization was performed on fixed 1-3 dpf zebrafish embryos using previously described protocol [18]. The stained embryos were imaged using a stereomicroscope (Zeiss STEMI 508 (Zeiss Oberkocken, Germany)). A 550 bp fragment of *elovl2* cDNA was PCR amplified and cloned into a TOPO vector (Invitrogen). (Primers 5’ F AGGCAGTCATTTAGGTGACACTATAGATGG and 3’ R CGTCGTGGACTAATACGACTCACTATAGAC). The plasmid was linearized and an *in vitro* transcription reaction with DIG labeling mix (Roche, cat. no. 1277073) was performed as previously described [18]. For staining of retinal sections, the RNAscope protocol was followed as per manufacturers protocol (ACD (Newark, CA); *elovl2* probe Cat. No. 550301). The optional step of target retrieval was excluded to allow better retention of the samples on the slides. The stained sections were flooded with Prolong^TM^ Gold Antifade Mountant (Cat. No. P36930) and covered with coverslips (Fisher Scientific. Waltham MA **12-545F**) before imaging.

### Microscopic Imaging

Fluorescent imaging of the fluorescent *in situ* hybridization (RNAscope) as well as the retinal anatomy analysis were performed using the Keyence microscope.

### Zebrafish visual motor response

Larvae at 6 dpf were placed in flat bottom clear 96 well plates (Corning CLS3370 Corning NY) and the assay was performed in DanioVision observation chamber (Noldus Information Technology, Wageningen, the Netherlands). The light program was designed based on a previously published protocol [19]. The larvae were dark-adapted for 3 hours, followed by three cycles of light on/off program. In each light cycle, white light at 100% intensity was turned on for 5 minutes followed by 30 minutes dark period. The activity of larvae was measured 30s before and after light on/off using EthoVision XT software (Noldus Information Technology, Wageningen, the Netherlands). A 5-minute dark period was included in the light program to measure the activity of larvae 30s before the first light on. The activity data was exported from the software and analyzed using Microsoft Excel.

### Zebrafish retinal anatomy analysis

The embryos were fixed in 4% PFA for 72 hours and transferred into 100% ethanol solution. The samples were submitted to HistoWiz (https://home.histowiz.com) for sectioning and H&E staining for histological analysis. For uniformity, only sections close to nerve fibers were selected for the measurements. Measurements were taken at three different regions in each selected section: near optic nerve (middle), anterior to the optic nerve (anterior) and posterior to optic nerve (posterior). Measurements were recorded using Fiji ImageJ software version 1.52p (National Institutes of Health, USA).

### Lipid analysis

For lipidomics sample preparation, 2 days and 7 days post fertilization (dpf) larvae were transferred in Eppendorf tube and excess water was removed. The tubes were then placed in a dish containing dry ice and ethanol to flash freeze the larvae. 12 embryos were placed in each tube, with 3 replicates per timepoint. Lipids were extracted using a modified version of the Bligh-Dyer method [20]. Briefly, zebrafish embryos were homogenized in 1 mL PBS and shaken in a glass vial (VWR) with 1 mL methanol and 2 mL chloroform containing internal standards (^13^C16-palmitic acid, d7-Cholesterol) for 30s. The resulting mixture was vortexed for 15s and centrifuged at 2400 x g for 6 min to induce phase separation. The organic (bottom) layer was retrieved using a Pasteur pipette, dried under a gentle stream of nitrogen, and reconstituted in 2:1 chloroform:methanol for LC/MS analysis.

Extracted lipids were resuspended in 200 μL of EtOH, incubated with 0.1 M KOH at room temperature for 24 hours for saponification. The reaction was stopped by addition of 0.2 M HCl. Lipids were extracted as described above with d31-palmitic acid as internal standard.

Untargeted lipidomic analysis was performed on a Vanquish HPLC online with a Q-Exactive quadrupole-orbitrap mass spectrometer equipped with an electrospray ion source (Thermo). A Bio-Bond C4 column (Dikma, 5 μm, 4.6 mm × 50 mm) was used. Solvent A consisted of 95:5 water:methanol, Solvent B was 60:35:5 isopropanol:methanol:water. Solvents A and B contained 5 mM ammonium formate with 0.1% formic acid. The gradient was held at 0% B between 0 and 5 min, raised to 20% B at 5.1 min, increased linearly from 20% to 100% B between 5.1 and 55 min, held at 100% B between 55 min and 63 min, returned to 0% B at 63.1 min, and held at 0% B until 70 min. Flow rate was 0.1 mL/min from 0 to 5 min, 0.4 mL/min between 5.1 min and 55 min, and 0.5 mL/min between 55 min and 70 min. Data was acquired in negative ionization mode. Spray voltage was −2.5 kV. Sheath, auxiliary, and sweep gases were 53, 14 and 3 arbitrary units (a.u.), respectively. Capillary temperature was 275°C. Data was collected in full MS/dd-MS2 (top 5). Full MS was acquired from 100–1500 m/z with resolution of 70,000, AGC target of 1×10^6^ and a maximum injection time of 100 ms. MS2 was acquired with resolution of 17,500, a fixed first mass of 50 m/z, AGC target of 1×10^5^ and a maximum injection time of 200 ms. Stepped normalized collision energies were 20, 30 and 40%.

Targeted lipidomic analysis was performed on a Dionex Ultimate 3000 LC system (Thermo) coupled to a TSQ Quantiva mass spectrometer (Thermo). A XBridge C8 column (Waters, 5 μm, 4.6 mm × 50 mm) was used. The solvents and gradient were as described above. MS analyses were performed using electrospray ionization in negative mode, with spay voltages of −2.5 kV, ion transfer tube temperature of 325 °C, and vaporizer temperature of 200 °C. Pseudo multiple reaction monitoring (MRM) was performed to detect fatty acids.

Lipid identification was performed with LipidSearch (Thermo). Mass accuracy, chromatography and peak integration of all LipidSearch-identified lipids were verified with Skyline [21]. Peak integration of targeted fatty acids was also performed with Skyline. Peak areas were used in data reporting, data was normalized using internal standards. The relative abundance of lipid classes were calculated by the percent relative area method with proper normalization using internal standard and considering the sum of all relative areas of the identified lipids.

Statistical analyses were conducted using Prism 7 (GraphPad Prism, La Jolla, CA, USA). All values are expressed as means ± SD (standard deviations). One-way ANOVA was performed to determine significant differences between different groups. Significant calls were made based on P values <; 0.05 and the |Fold Change (FC)| >1.5.

## RESULTS

### Zebrafish *ELOVL2* is well conserved compared to other higher vertebrates

We first investigated the conservation of zebrafish ELOVL2 protein to other vertebrate species (Supplementary Figure 1A). There is a single zebrafish *elovl2* ortholog based on BLAST searches. We performed protein sequence alignment between zebrafish and other vertebrate species (Supplementary Figure 1A, B). Multiple sequence alignment of ELOVL2 proteins from these species showed high sequence conservation in the transmembrane regions (Supplementary Figure 1B, red highlight). In total, the zebrafish ELOVL2 protein has 65.2% sequence conservation to the human Elovl2 protein. Of note, critical residues such as human amino acid 234 cysteine is well conserved (Supplementary Figure 1B, green highlight) which provides substrate specificity of the enzyme [8].

### *Elovl2* is highly expressed in Zebrafish retina

Next, we investigated the expression of *elovl2* in both zebrafish embryonic development as well as adult, focusing on the eye. Strong expression of *elovl2* was observed in the eye as well as in the hindbrain at day 3 (Figure 1C, C’).

**Figure 1.**
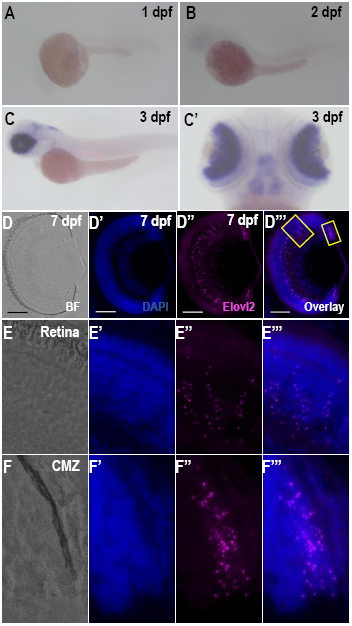
A-C Expression of elovl2 in zebrafish larvae. Lateral view of elovl2 staining in (**A**) 1 dpf and (**B**) 2 dpf zebrafish retinas. Lateral (**C**) and dorsal (C’) view of elovl2 staining in 3 dpf zebrafish retina, respectively. B-G. RNAscope in situ hybridization with elovl2 probe in 7 dpf zebrafish retina showing brightfield (**D, E, F**), DAPI (D’, E’, F’), elovl2 (D’’, E’’, F’’) and overlay (D’’’, E’’’, F’’’) images. Scale bars = 50μm. Magnified view of boxed regions in Figure 1D’’’. The magnified views of Retina (Panel E) and CMZ (Panel F) showing *elovl2* staining in 7 dpf zebrafish retina.

Using RNAscope *in situ* hybridization on fixed retinal sections, *elovl2* expression was studied in developing (7 dpf), young adult (3 mpf) and old (14 mpf) zebrafish retinas. The 7dpf retina showed the significant expression of *elovl2* at the ciliary margin region (Figure 1D’’ and 1D’’’, small rectangle and Supplementary Figure 2A’’’). In the other regions of the retina, the punctate RNAscope signal of *elovl2* probe was detected across all the retinal layers with the particular enrichment in the inner nuclear layer (INL) (Figure 1D’’’, larger boxed area and Supplementary Figure 2B). To determine the changes in *elovl2* expression with age, we examined the expression of *elovl2* in 3 mpf and 14 mpf adult zebrafish retinas (Supplementary Figure 2). Similarly, to observations at 7dpf, higher expression of *elovl2* was observed in the INL compared to the other retinal layers at adult stages (Supplementary Figure 2). Interestingly, although the older retina at 14 mpf also showed higher expression of *elovl2* in the INL compared to other layers, an overall reduction in *elovl2* expression in 14-month zebrafish retina compared to 3-month zebrafish retinas was observed (Supplementary Figure 2).

**Figure 2.**
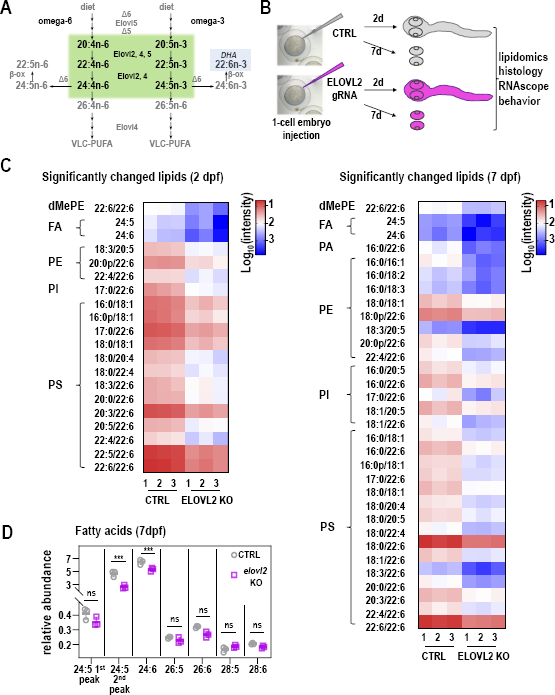
Generation and analysis of *elovl2* crispants. **A**. ω-3 and ω-6 PUFA Elongation pathway. **B**. Workflow for creating biallelic *elovl2* mutants (‘crispants’) in zebrafish using CRISPR/Cas. **C**. Significantly changed lipids in *elovl2* crispants at day 2 and day 7. **D**. Relative abundance of PUFAs in control and *elovl2* crispants shows lower abundance of DHA.

### Elovl2 crispants disrupt elovl2 coding sequences and have functional effects on fatty acid elongation

To determine the function of *elovl2* during zebrafish development, we generated biallelic *elovl2* mutants (‘crispants’) using CRISPR-Cas9 technology (Figure 2B). To knockdown *elovl2*, we injected gRNAs targeting exon 2 and 3 of the zebrafish *elovl2* gene (Supplementary Figure 3). Sequence analysis of the target region showed high efficiency of both gRNAs (Supplementary Figure 3).

DNA sanger sequencing demonstrated that over 85% of DNA was biallelically edited using the ICE v2 Crispr gene editing tool. Analysis of other loci did not show any gene editing (data not shown).

We then assessed whether *elovl2* crispants had any changes in fatty acids, particularly the Elovl2 substrates and direct products. Untargeted lipidomic analysis was performed on whole zebrafish embryo (day 2) and on embryos eyes (day7). Overall, 110 lipid species in 11 lipid classes (Supplementary Figure 4) were identified, including Free Fatty acids (FFA), Ceramides (Cer), Dimethylphosphatidylethanolamine (dMePE), Lysophosphatidylethanolamine (LPE), Phosphatidic acid (PA), Phosphatidylcholine (PC), Phosphatidylethanolamine (PE), Phosphatidylglycerol (PG), phosphatidylinositol (PI), phosphatidylmethanol (PMe) and phosphatidylserine (PS). The relative signal of each lipid class of control and *elovl2* crispants was analyzed (Supplementary Figure 5). FAs were the lipids which showed the highest signal in both groups on Day 2 and Day 7. On Day 2, significant increases in relative signal were observed for FA (29.8%; *P* = 0.0010), PC (35.4%, *P* = 0.0142), and PMe (24.4%, *P* = 0.0372) in *elovl2* crispants, dMePE, PE, PG, PI, PS were decreased by 80.0%, 33.3%, 46.4%, 49.4% and 52.9%, respectively. Day 7 *elovl2* crispants only showed a significant increase in FA, while Cer, dMePE, LPE, PA, PE, PG, PI, PMe and PS were significantly decreased in the extract. Further analysis showed downregulation of 20 and 32 lipids, respectively (p<;0.05, |log2(Fold change)| > 1.5)(Figure 2C), including dMePE, FA, PE, PI, PS at day 2, and dMePE, FA, PE, PI, PS at day 7. Interestingly, most significant decreases were observed for lipids containing at least one fatty acids tail of 20:4, 20:5, 22:4, 22:5, 22:6, 24:5 and 24:6, which are products of *elovl2* in LC-PUFAs pathway. Lastly, targeted analysis of very long chain fatty acids revealed significant loss of 24:5 and 24:6 fatty acids (Figure 2D), crucial substrates for VLC-PUFA synthesis. These data revealed that targeted elimination of *elovl2* significantly affects the biosynthesis of long and very long chain PUFAs.

### Elovl2 knockdown disrupts the thickness of retinal layers

To determine the effect of *elovl2* knockdown on the retinal architecture images of H&E stained retinal sections of *elovl2* and control crispants were analyzed. No gross changes in the retina morphology was observed (Figure 3A). However, when the thickness of the individual retinal layers of *elovl2* and control crispants was plotted (Supplementary Figure 6, Figure 3B), day 7 *elovl2* crispants showed an increase in the thickness of several retinal layers (Figure 3B). In particular, we found a significant increase in the thickness of RGL, IPL, INL, ONL, and PRL. Figure 3 and Table 1 shows the difference in the thickness observed in each of these layers at three different regions measured. Since the thickness of retinal layers vary along the retina, percent change in lengths of each layer showing significant difference in *elovl2* mutants compared to controls was calculated (Table 1). The increased thickness of retinal layers in cripants was found to be more profound at the region surrounding the optic nerve (Table 1, middle column). The retinal ganglion cell layer and inner nuclear layer were found to be most affected by *elovl2* knockdown. This data suggests that the *elovl2* mutation affects the mechanism responsible for the maintenance of retinal architecture.

**Figure 3.**
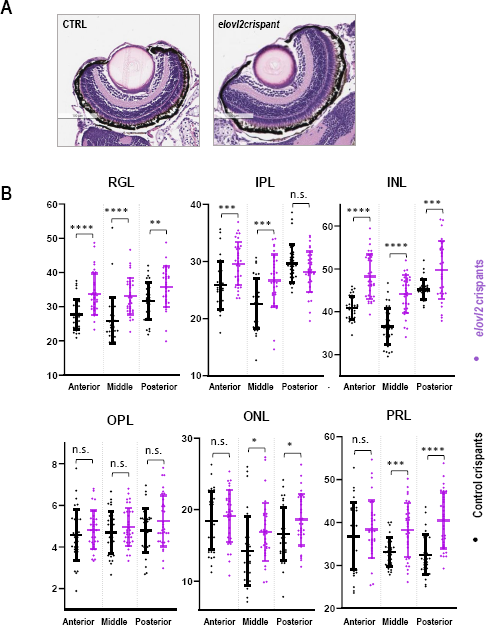
**A**. H&E stained retinal sections of *elovl2* and control crispants. **B**. The thickness of the individual retinal layers of *elovl2* and control crispants. Plots showing comparison of lengths of different layers in retina between control and *elovl2* crispants. The measurements were taken at three different regions in the retina termed as anterior, posterior and middle. RGL – Retinal Ganglion Cell Layer, IPL-Inner Plexiform Layer, INL-Inner Nuclear Layer, OPL-Outer Plexiform Layer, ONL-Outer Nuclear Layer, PRL-Photoreceptor Layer. (*P <; 0.05; **P <; 0.01; ***P <; 0.001; ****P <; 0.0001)

### Elovl2 is expressed in Muller glia

A previous study in zebrafish correlated increase in retinal layer thickness with lack of the Muller glia cells [22]. To verify whether in *elovl2* is expressed in Muller glia we have performed RNAscope experiment with a probe against glutamine synthetase (*glula),* a specific marker for Muller glia, and *elovl2* in adult zebrafish. As presented on Figure 4A, *elovl2* RNA was clearly detectable and overlapping with *glula* signal in the inner nuclear layer and inner plexiform layer. To investigate whether lack of elovl2 causes significant changes in Muller Glia abundance we have performed RNAscope experiment on day 7 wild-type and *elovl2* crispants embryonic eyes. Our data show (Figure 4B) that *glul* signal was easily detectable on the comparable levels in both genotypes suggesting that Muller glia are intact in crispants eyes.

**Figure 4.**
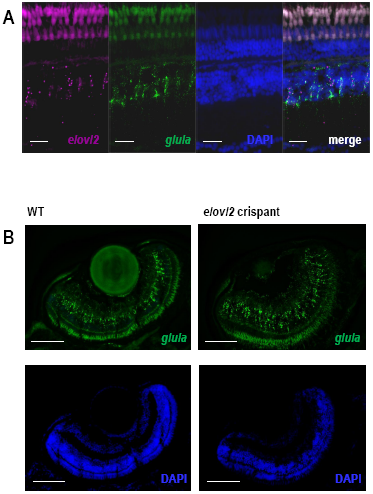
**A**. Expression of *elovl2* RNA and Muller Glia specific gene as detected by RNAscope in normal adult zebrafish. **B**. Expression of *glula* in 7dpf control and elovl2 crispants shows no changes of *glula* expression upon *elovl2* knockout.

### Elovl2 crispants show changes in visual behavior

To determine the effect of *elovl2* knockdown on visual function in zebrafish, we performed the motor response (VMR) assay at 6 dpf. The VMR is a well-studied visual behavior reflex which is a startle response in reaction to bright light (Figure 5A). This response begins to manifest on day 3 but becomes more robust by day 5 [23]. There are 2 elements to the VMR, the Light-on reflex (VMR-ON) which is a sharp increase in locomotor activity at the onset of light with a return to baseline activity in 30 seconds, and the Light-off VMR (VMR-OFF), which is a sudden increase in locomotor activity at the light offset, which then gradually returns to baseline over 30 minutes [24].

**Figure 5.**
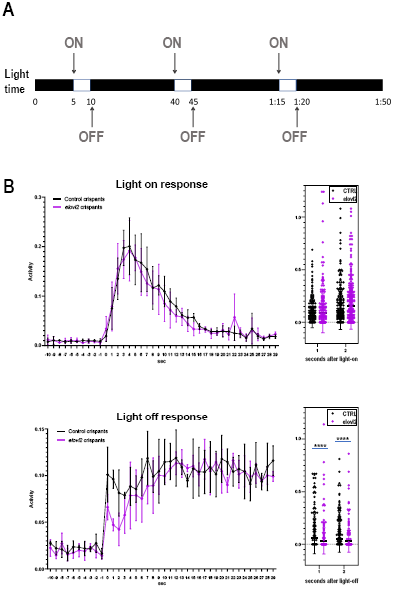
Zebrafish lacking elovl2 activity show visual phenotype. **A**. Diagram of motor response (VMR) assay. (B) Motor response of control and elovl2 crispants to light-on and light-off stimulus. **B**. The thickness of RGL, IPL, INL, ONL, and PRL at three different regions.

The control and *elovl2* crispants showed exponential increase in the activity in response to light-on stimulus (Figure 5B). The activity peaked between four to six seconds after light on stimulus in both groups. However, we did not observe any significant difference between control and *elovl2* crispants for VMR-ON assay. Both groups showed gradual decrease in mean activity of larvae to baseline between 15-30 seconds. Therefore, we concluded that the *elovl2* knockdown does not affect the VMR-ON response in zebrafish at 6 dpf.

We further analyzed the activity of larvae in response to light-off stimulus. We observed the sharp VMR-OFF response in control and *elovl2* larvae (Figure 5B and Supplementary Figure 7). The activity of the larvae first peaked around 1-2 seconds after light-off stimulus followed by short decline. Unlike light-on stimulus, the activity of larvae was peaked again after short decline (Figure 5B). We observed the VMR-OFF response was ~50% weaker than VMR-ON within same group of larvae (Supplementary Figure 7, control crispants: panel A vs *elovl2* crispants: panel B). Interestingly, the *elovl2* crispants showed much weaker activity in response to light-off stimulus as compared to control crispants. We calculated the activity difference between control and *elovl2* crispants at one and two seconds after light-off stimulus. The *elovl2* crispants showed 48.96% (P = 0.0004) and 51.6% (P = 0.0007) less activity compared to control crispants at one and two seconds after light on, respectively (control crispants, n=372; *elovl2* crispants, n = 336).

## DISCUSSION

We have recently implicated ELOVL2 as a molecular regulator of aging in the mouse retina [11]. In this work, we investigated the expression and function of *elovl2* in the embryonic zebrafish retina. We found that *elovl2* is strongly expressed in the zebrafish retina during early embryogenesis as well as in the adult retina, with expression primarily confined to Müller glia. Analysis of *elovl2* crispants results in changes in lipid levels in the zebrafish embryo in particular lipids containing polyunsaturated fatty acids longer than 22 carbons. Analysis of *elovl2* crispants demonstrates no gross changes in retinal morphology, but significant changes in the VMR-OFF response further underlying importance of the enzyme in the healthy vision.

The *elovl2* enzyme is strongly conserved between zebrafish and other species. Interestingly, while work has shown that zebrafish *elovl2*, elongates C20 and C22 carbons, it also has some elongation activity for C18 fatty acids which is different from mammalian *Elovl2* (10,11). This is likely due to some overlap of function between *elovl2* and *elovl5* in zebrafish (Figure 2A), however recent work suggests that *elovl2* is the primary elongase required for the synthesis of DHA [25]. Interestingly, the zebrafish *elovl4* enzyme is also able to elongate C20 and C22 fatty acids in addition to the activity towards VLC-PUFAs.

When investigating the expression of the *elovl2*, we observed a strong expression at 3 dpf. On histological sections at 7 days, clear RNAscope signal can be observed throughout the retina. *In situ* hybridization of retina sections in adult zebrafish showed similar pattern of the expression. A lower expression of *elovl2* in old zebrafish compared to young adult zebrafish was observed, suggesting that there may also be an age dependent decrease in *elovl2* expression that has been seen in human and mouse tissues [9]. Intriguingly, we find expression of *elovl2* primarily in the Müller glia, which is in contrast to the mouse data where it is primarily expressed in photoreceptors in the retinal pigment epithelium.

Several groups analyzed expression patterns of *elovl2, 4* and *5* in zebrafish retina[14, 16, 26]. Our data is in general agreement with published data, although one of the groups detected the first expression of Elovl2 earlier than at 3dpf [26] Intriguingly none of the studies convincingly presented the expression of *elovl5* in the eye while *elovl4* seem to be expressed mostly in retinal pigmented epithelium cells in striking contrast to the mammalian gene. Taken together the data suggests that the *elovl2* is the sole fatty acid elongase able to elongate C22 to C24 in inner nuclear layer.

We then assessed the function of Elovl2 in zebrafish embryos by creating *elovl2* crispants. This was done by injecting 2 gRNAs specific to *elovl2* as well as Cas9 protein into single cell embryos. The two gRNAs when injected in tandem, high rates of gene editing was observed (Supplementary Figure 3), causing biallelic changes in *elovl2* gene. No off-target effect has been observed.

*Elovl2* crispants had significant changes in levels of various fatty acids, consistent with loss of Elovl2 function compared to control crispants. Lipidomics performed on whole zebrafish embryo showed a significant decrease of LC-PUFAs in *elovl2* crispants. These FAs, including 20:4, 20:5, 22:4, 22:5, 22:6, 24:5 and 24:6, are potential products of Elovl2 enzymatic activity. In addition, the availability of specific fatty acids affected the overall composition of lipids in the zebrafish embryos. In particular levels of phospholipids were significantly changed while availability of free fatty acids substantially increased. This data shows that lack of specific fatty acids can affect the lipid biosynthesis, therefore the overall compositions of membranes.

To analyze the impact of the lack of *elovl2* enzymatic activity on the retinal morphology thickness of the retinal layers in the 7dpf control and *elovl2* crispants sections was examined and quantified. No gross disruptions of the retinal architecture were observed, however, the thickening of several retinal layers in *elovl2* crispants when compared to controls was detected. Other studies have reported changes in the thickness of retinal layers after the inactivation of Müller glia cells [22, 27]. MacDonald et. al. showed that inhibition of Müller glia formation during zebrafish development results into wider RGL but no effect was observed in other retinal layers. This suggest that other factors play role in maintenance of retinal layers in absence of Müller glia. While our observation further supports the indispensable role of Müller glia in maintenance of RGL in zebrafish retina, the factors playing involved in the maintenance of other retinal layers in the absence of Müller glia cells remain unstudied. Another study targeting the deletion of Dicer1 miRNA specifically in Müller glia cells present in INL of mice retina reported increase in thickness of INL. Intriguingly, the group also observed reduction in thickness of ONL. Considering the global change in the membrane lipids composition in *elovl2* crispants (Supplementary Figure 5) and therefore different physical properties of the membranes, our studies further supports the observation of the key role of the Müller glia in the maintenance of retinal architecture.

In the mammalian retina, *Elovl2* is expressed in photoreceptors, especially in cones. Since the VLC-PUFAs are the key fatty acids in photoreceptor disks membranes, the striking difference in the expression pattern of mammalian and zebrafish *elovl2* gene suggests a potential dependence of photoreceptor cells on Muller glia. This interesting interaction would involve the Muller glia in zebrafish eye to provide VLC-PUFAs for the photoreceptor disks during the daily disk regeneration. Further studies are required to investigate this hypothesis.

Finally, we examined the changes in visual behavior in zebrafish *elovl2* crispants. We performed the VMR assay, a well characterized visual reflex that develops by day 5. We assessed the visual motor response at day 6 and did not find any changes in the VMR-ON response. However, we did see a significantly weaker, albeit still present, VMR-OFF response after *elovl2* knockdown. It is still unclear what parts of the visual circuit mediate this VMR-OFF response. Interesting, *slc7a14* mutant zebrafish, which affects rod photoreceptors more than cones, shows greater defects in the VMR-OFF response than the VMR-ON response [28]. This is in contrast to *Pde6c* mutation, which results in cone degeneration, and leads to defects in the VMR-ON response rather than VMR-OFF response [19]. This suggests a possibility that VMR-ON and VMR-OFF responses may be primarily influenced by different classes of photoreceptors [28]. Further studies are needed to examine the effect of *elovl2* on visual function in the zebrafish retina.

In conclusion, this is the first study to investigate the role of *elovl2* in the zebrafish retina. Loss of *elovl2* activity results in broad changes across the production of lipids compatible with its function as an elongase of LC-PUFAs as well as in impact on visual function. *Elovl2* expression in the Müller glia may suggest additional and unique functions of zebrafish *elovl2* compared to mammals. Taken together, our work further advances our understanding of the role of *elovl2* and LC-PUFAs in retina structure and function.

## Acknowledgement

We thank Michal Krawczyk for help with preparing the figures. This work was funded by an unrestricted Research to Prevent Blindness grant to Gavin Herbert Eye Institute at University of California, Irvine, an unrestricted Research to Prevent Blindness grant to University of California, San Diego, Research to Prevent Blindness Special Scholar award to D.S.K. and NEI K08EY030510 to D.L.C., as well as a Faculty senate grant from University of California San Diego to D.L.C.

## Contributions

M.D. acquiring experimental data, analysis of data, and writing of the manuscript.

F.G., Q.X acquiring experimental data, analysis of data, and writing of manuscript

D.V.F, E.Z., A.F.M.P. A.S. acquiring experimental data and analysis

D.S.K. D.L.C, conceptualization of experiments, writing and editing of manuscript

---

**Supplementary Figure 1.**
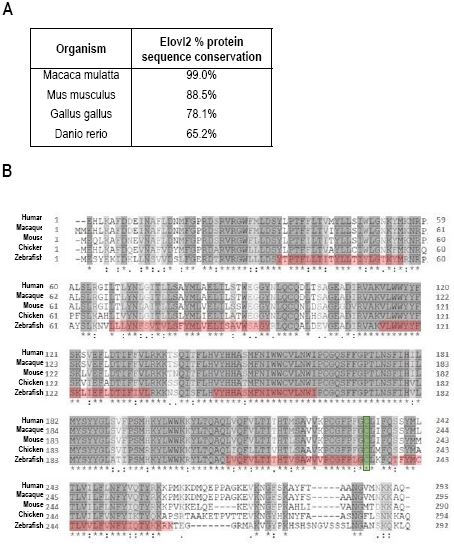
**A**. Conservation of *elovl2* gene and protein sequence across different species. **B**. Multiple sequence alignment of ELOVL2 protein in different species showing conserved sequences. Amino acids forming transmembrane helices are highlighted in red; * represents identical amino acids;: represents amino acids with highly similar properties;. represents amino acids with weakly similar properties.

**Supplementary Figure 2.**
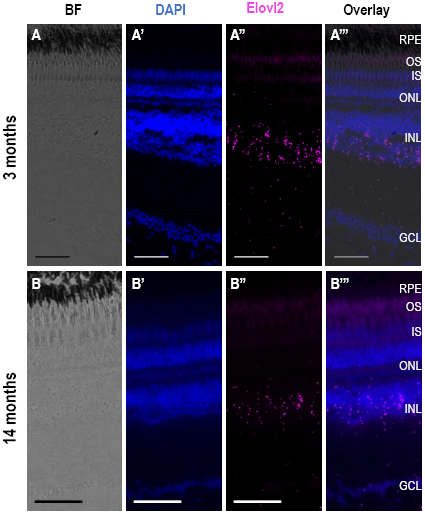
Expression of *elovl2* in adult zebrafish retina. Expression analysis of elovl2 in 3 months (Panel A) and 14 months (Panel B) old zebrafish retina showing brightfield (A, B), DAPI stained (A’, B’), elovl2 stained (A’’, B’’) and overlay (A’’’, B’’’) images. Scale bars = 50 μm.

**Supplementary Figure 3.**
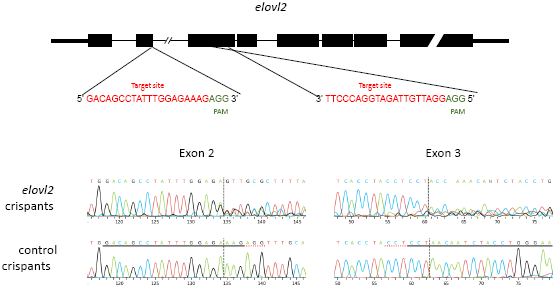
Sequence analysis of elovl2 and control crispants at exon 2 and exon 3.

**Supplementary Figure 4.**
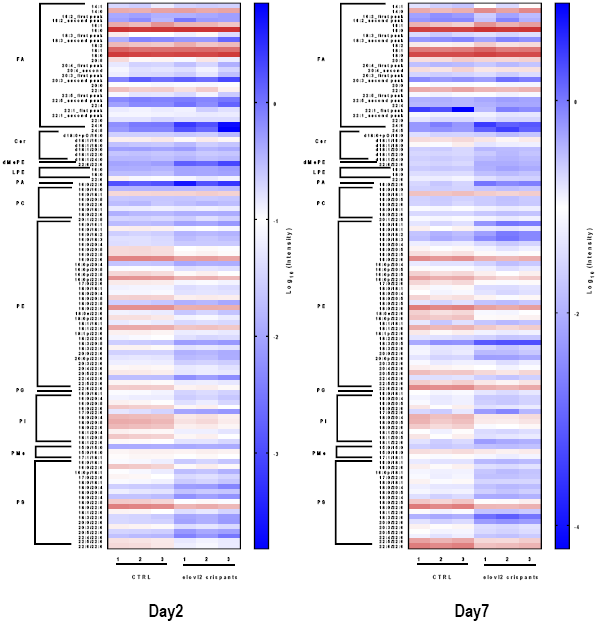
Heatmap of all the identified lipids on whole zebrafish embryo (day 2) and on embryos eyes (day7)

**Supplementary Figure 5.**
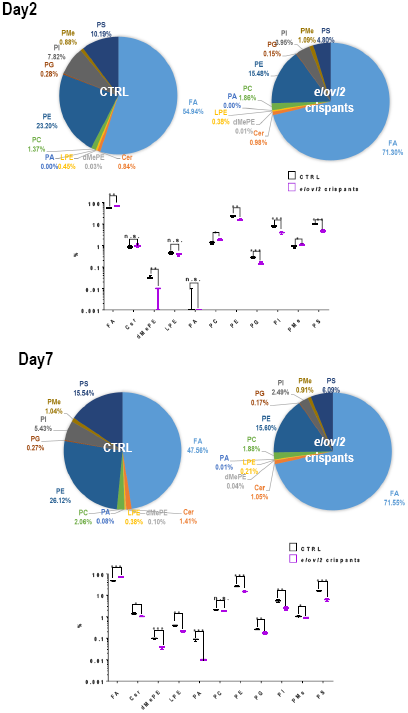
The relative abundance of each lipid class of control and elovl2 crispants

**Supplementary Figure 6.**
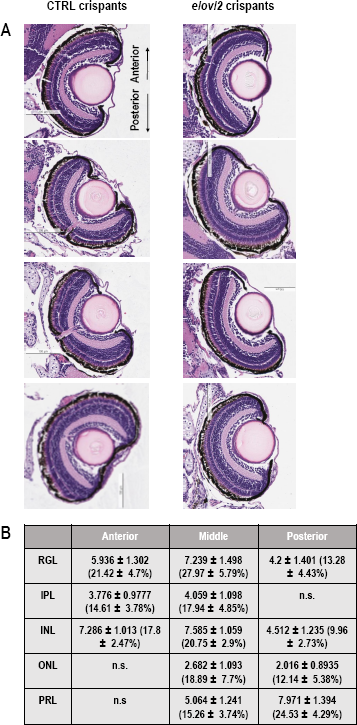
Representative H&E staining images of elovl2 and control crispants showing different layers of retina. Scale bars = 100 μm.

**Supplementary Figure 7.**
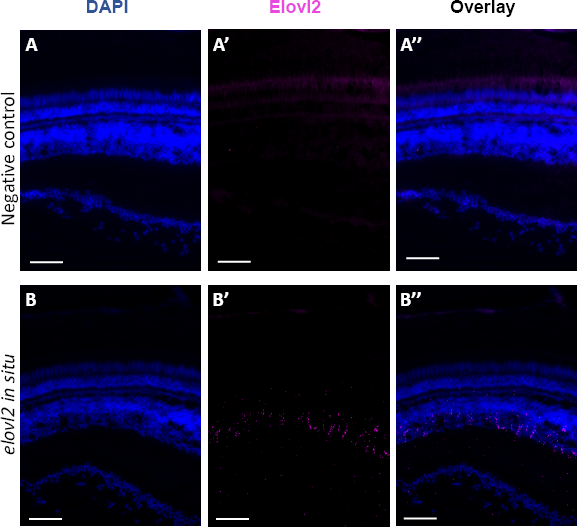
Comparison of staining intensity with *elovl2* and control probe. The negative control probe (Panel **A**) showed light autofluorescence in photoreceptor layer (A’) confirming specific staining of elovl2 (B’) in the 3 months old retina (Panel **B**). Scale bars = 50 μm.

